# Source identity shapes spatial preference in primary auditory cortex during active navigation

**DOI:** 10.1101/2021.01.05.425444

**Authors:** Diana Amaro, Dardo N. Ferreiro, Benedikt Grothe, Michael Pecka

## Abstract

Localizing and identifying sensory objects while navigating the environment are fundamental brain functions. However, how individual objects are neuronally represented during unrestricted self-motion is mostly unexplored. We trained gerbils on a behavioral foraging paradigm that required localization and identification of sound-sources during free navigation. Chronic multi-electrode recordings in primary auditory cortex during task performance revealed previously unreported sensory object representations. Strikingly, the egocentric angle preference of the majority of spatially sensitive neurons changed significantly depending on the task-specific identity (outcome association) of the sound-source. Spatial tuning also exhibited larger temporal complexity. Moreover, we encountered egocentrically untuned neurons whose response magnitude differed between source identities. Using a neural network decoder we show that together, these neuronal response ensembles provide spatio-temporally co-existent information about both the egocentric location and the identity of individual sensory objects during self-motion, revealing a novel cortical computation principle for naturalistic sensing.

**Highlights:**

- Localization task during free navigation prompts diverse spatial tuning in gerbil A1
- Spatial preference of individual neurons changes with sound-source identity
- Ego- and allocentric information are spatio-temporally coexistent in A1 ensembles
- Active sensing reveals new cortical representations for sensory object identification

## Introduction

Information about the position of sensory objects and identifying their concurrent behavioral relevance is vital to navigate the environment. In the auditory system, neurons compute azimuthal spatial information via angle-specific differences of sound features between the ears. From brainstem to primary auditory cortex (A1), a predominance of broad neuronal tuning to contralateral sound-source locations has been reported, with a smaller subset tuned to ipsilateral or frontal positions (Grothe et al., 2010; Mickey and Middlebrooks, 2003; Mrsic-Flogel et al., 2005; Stecker et al., 2005; Woods et al., 2006). Accordingly, spatial tuning is assumed to be egocentric (i.e., based on the position of the sound-source relative to the observer). Nevertheless, most studies on spatial tuning to date were conducted in either anesthetized or head-fixed and passively listening animals, thus lacking important aspects of real-life localization behavior that may crucially impact the nature of spatial coding and sensory object representation (van der Heijden et al., 2019).

Movements and active exploration are fundamental components of natural sensing (Schroeder et al., 2010). Notably, while self-movement constantly alters the egocentric sound-source location, the perception of source position remains stable relative to the world coordinates, i.e. is allocentric (Burgess, 2008). Recent results from one study with freely moving ferrets suggest the existence of allocentric representation in a small minority of neurons in A1 (Town et al., 2017). However, the stimuli and associated sources in this study were task-irrelevant since subjects were passively exposed to sounds while searching for water. Yet active sensing and task engagement / stimulus relevance critically modulate neuronal coding in A1 (Benson et al., 1981; Fritz et al., 2003; David et al., 2012; Lin et al., 2019; von Trapp et al., 2016; Lee and Middlebrooks, 2011; Kato et al., 2015), and thus it remains unclear how sound-sources are represented during unrestricted movement and active localization. Here, we took advantage of the recently developed Sensory Island Task (SIT) paradigm (Ferreiro et al., 2020) and recorded from A1 neurons of freely exploring animals that actively localized sounds from sources with distinct task identities (i.e., associated behavioral outcome) and allocentric locations (Figure 1A). We find that the spatial tuning in A1 during active sensing deviated profoundly from the canonically assumed egocentric representation. The majority of neurons exhibited temporally diverse spatial tuning that differed between sound-sources. Artificial neural network decoding demonstrates that on the population level these novel tuning features generate spatio-temporally coexistent information about the instantaneous source-angles (egocentric reference frame) and angle-independent source identity (allocentric reference frame).

**Figure 1.**
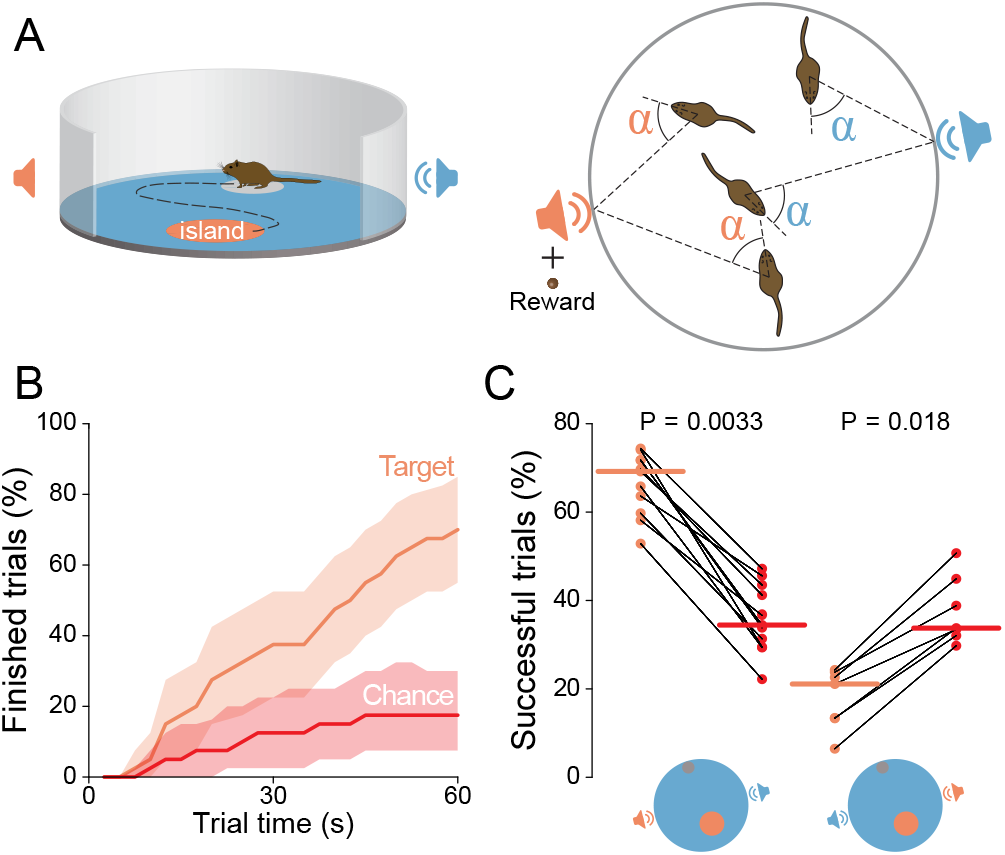
Sound localization task. (A) Left: general scheme of the task: the animal starts a trial in the initiation platform and hears a pulsed harmonic complex from the background loudspeaker until it enters the island, which triggers a switch to the target loudspeaker; to report the switch the animal must stay within the island for 6 seconds. Right: Egocentric sound-source angles are ambiguous and hence insufficient for task completion. Pairing of the target loudspeaker with reward delivery allows the formation of source-specific identity associations. (B) Task performance of one animal in comparison to chance during a particular session as a function of time spent in the trial (40 trials, shaded areas correspond to 95% confidence interval). (C) Left: task performance at maximum trial length (60 s) in comparison to chance (11 animals, Wilcoxon signed-rank test, horizontal lines represent the median, previously reported – Ferreiro et al., 2020). Right: task performance at maximum trial length (60s) in comparison to chance in catch-trials with reversal of the reward contingency (5 animals, Wilcoxon signed-rank test, horizontal lines represent the median, previously reported – Ferreiro et al., 2020).

## Results

To investigate potentially undiscovered spatial representations during active sensing, we trained Mongolian gerbils *(Meriones unguiculatus*) to report whenever the presentation of a pulsed harmonic stack switched from a “background” loudspeaker to a “target” loudspeaker (separated by 180° in a circular arena, Figure 1A). The source change was triggered whenever gerbils entered a specific target area in the arena (the “island”), whose position was randomized across trials. The animals were trained to report the detection of this change by remaining within the island for 6 seconds to receive a food reward. Thus, activity of the target loudspeaker was associated with imminent reward delivery. The sounds emitted by the two loudspeakers were identical, hence completion was achievable only by determining the location of the active sound-source. Importantly, because the egocentric sound-source location can be the same for both loudspeakers, this information was rendered insufficient for task completion (Figure 1A and Video S1). Nonetheless, the animals reliably exhibited highly significant performance levels in reporting the activity of the target loudspeaker (Figures 1B and 1C), their behavior demonstrates allocentric localization (e.g. “loudspeaker to the west of the initiation platform”). Note that switching the rewarded loudspeaker identity in catch-trials resulted in performances significantly below chance level (Figure 1C), corroborating a task-specific identification of the allocentric location of loudspeakers by the animals. Performance remained similar before (mean success rate above chance level ± 95% confidence interval: 19.7 ± 5.7%) and after (20.2 ± 5.1%) physically swapping the loudspeakers (but not their task-specific identity), showing that animals were not using potentially existing specific spectral cues from the loudspeakers to solve the task. This allowed studying how listening to two sound-sources with distinct behaviorally meaning that are defined by their world-based position influences the spatial representation in A1.

To investigate the neuronal processing during task performance, expert animals were implanted with custom-made tetrodes to allow chronic recordings of action potentials from neurons in A1 (Figure 2A). Overall, we acquired responses during task performance from 364 single neurons and 246 multi-units from 5 gerbils. Activity in A1 was strongly correlated with the subjects’ goal-specific behavior, as response rates to the target loudspeaker were significantly lower when animals correctly finished the trial as opposed to when they wrongly left the island (Figures 2B and S1).

**Figure 2.**
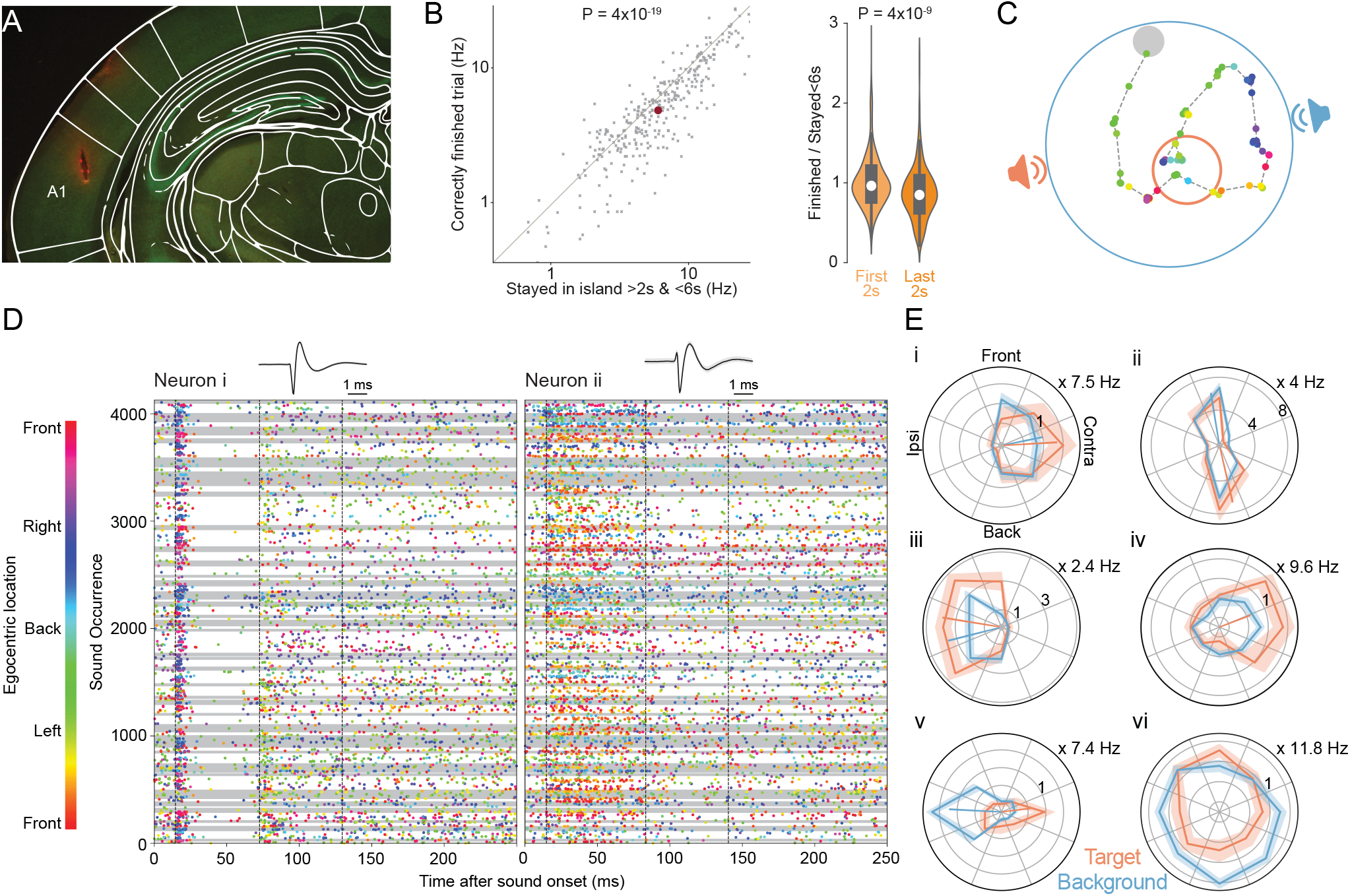
Non-canonical spatial tuning. (A) Reconstruction of tetrode position from histological information, superimposed with the location of A1 (Radtke-Schuller et al., 2016). (B) Left: Firing rates were significantly lower in the last 2 s when the animal finished a trial correctly compared to when it wrongly abandons the island (321 neurons, Wilcoxon signed-rank test, mean firing rate). Right: This decrease in firing rate was enhanced with more time spent in the island (Mann-Whitney U-test, first 2s: 346 neurons, last 2s: 321 neurons). Violin plots surrounding boxplots (the median is represented by a white dot) depict the distribution of the ratios between the mean firing rate when the animal correctly finished a trial and when it wrongly left the island. (C) Trajectory of the animal during a successful trial. Colored dots indicate the position of the animal at the moment of sound presentation and corresponding egocentric locations of the active loudspeaker. The color code is the same as in (D). (D) Raster plots of neuronal activity in two A1 neurons during task performance in the session represented in Figure 1B. Periods of stimulation by the target loudspeaker are highlighted by gray areas. Egocentric location of the active loudspeaker at the moment of the sound occurrence is color-coded. (E) Spatial tuning for six representative neurons for the target and background loudspeakers. The shaded area corresponds to the 95% confidence interval and the colored lines indicate the preferred tuning angle, with the length scaling with the vector strength. Neurons i and ii correspond to the ones depicted in (D).

Neuronal firing rates exhibited significant modulation (see below and Methods) as a function of the active loudspeaker’s angle relative to the animal’s body-axis (Figure 2C, see Methods) at specific time periods relative to sound presentation (Figure 2D, “onset period”: from the determined response latency + stimulus duration of 57 ms, 118 neurons; “offset period”: end of onset period + 57 ms, 58 neurons; “late response period”: remaining time between the periodic response to the stimuli, 42 neurons, see Methods). Construction of polar plots from these responses displayed a large variety of egocentric spatial sensitivity (Figure 2E). Alongside a fraction of “canonical” neurons that were similarly tuned to either loudspeaker with contralateral or ipsilateral preference (“i” and “iii” in Figure 2E; onset period: 37/118 = 31.3%; offset period: 7/58; late response period: 0/42), we observed neurons whose tuning to either loudspeaker can be described as orientation-sensitive (onset period: 16/118 = 13.6%; offset period: 1/58; late period: 2/42), with an apparent bias to the front/back orientation (“ii” in Figure 2E, comparable to reports in awake primates - Remington and Wang, 2019).

Remarkably, the spatial tuning of a large fraction of the neurons was source-sensitive, as their responses differed for the two loudspeakers (Figures 2E, 3A, and S2A). These neurons were either only significantly spatially tuned to one of the two sound-sources (“iv” in Figures 2E and 3A; 47/65 = 72.3 %) or exhibited a large tuning difference between the two sources (difference in preferred egocentric angle > 90°, “v” in Figures 2E and 3A; 18/65 = 27.7%, see Methods). This variety in tuning types was evident during all three response epochs inclusively during the onset period (and in cells with short onset latencies, Figure S2B), and particularly prominent during the late response period (onset period: 65/118 = 55.1%; offset response: 46/58 = 79.3%; late response period: 41/42=97.6%, Figure S2A). Source-specific tuning also occurred on the level of multi-unit responses (Figure S2C). Quantification of spatial selectivity exposed population-wide source-specific tuning, as vector strength (see Methods) was significantly higher for the target loudspeaker (Figure 3B).

**Figure 3:**
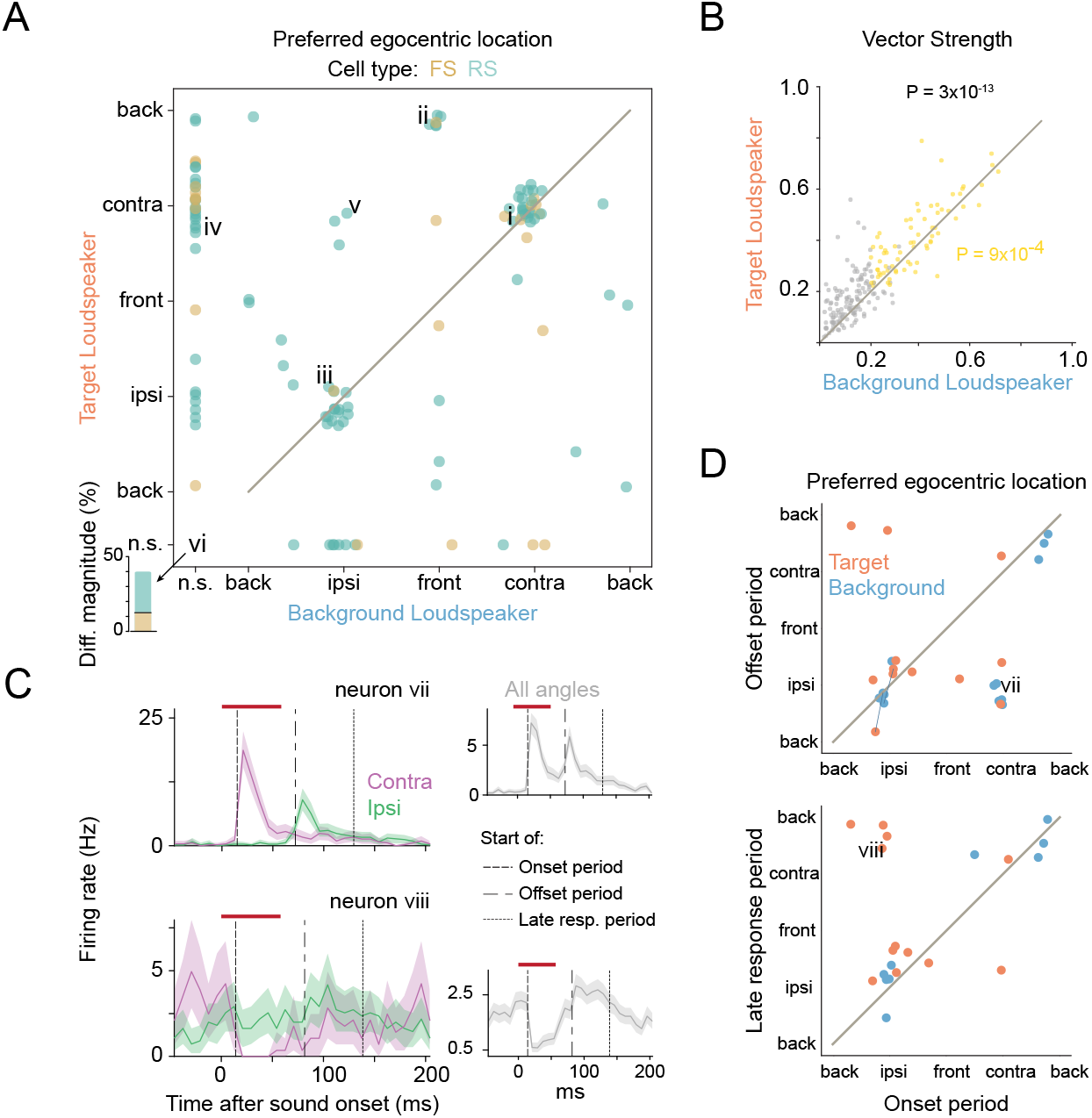
Complexity of neuronal responses in a context-dependent manner. (A) Correspondence of preferred egocentric locations between the target and the background loudspeakers. Color-code depicts putative neuronal type; FS: fast-spiking, RS: regular spiking (118 neurons, 71 of which tuned for both loudspeakers). Inset in lower left corner shows percentage of neurons that were not significantly (n.s.) tuned to either loudspeaker, but exhibited a significant difference in overall response magnitude (16/40 neurons, 5FS, 11 RS). (B) Vector strength was larger for the target than the background loudspeaker (Wilcoxon signed-rank test). Yellow data points represent neurons spatially tuned to both loudspeakers (71 neurons) and data from neurons that were not significantly spatially modulated for at least one of the loudspeakers are shown in gray (133 neurons). (C) Peristimulus time histograms (PSTH) for two representative neurons with temporally complex spatial tunings. The shaded area corresponds to the 95% confidence interval, the red line represents the sound stimulation period and the vertical lines indicate the time period considered for the construction of the spatial tuning (i.e. onset, offset and late response periods). The violet/green PSTHs correspond to responses acquired only for contra- or ipsilateral loudspeaker locations (in a 90^°^ angular bin). (D) Spatial tuning comparison between the onset and the offset time periods (upper panel, 23 neurons) and between the onset and the late response periods (lower panel, 21 neurons). The dark blue lines connect background and target preferred egocentric locations that correspond to the same neuron.

So far, our data suggests that active localization during self-motion in SIT reveals a variety of previously unreported characteristics of spatial sensitivity. However, movement can modulate cortical activity unspecifically (Zhou et al., 2014; Schneider et al., 2014). We therefore performed additional control analyses with data restricted to neuronal responses from periods when the animals were not moving (no translational or rotatory body movements). This resulted in qualitatively similar observations (Figure S2D), demonstrating that our findings are not caused by unspecific movement modulation.

A dependency of tuning preferences on the putative neuronal type was not observed (regular or fast spiking, Figures 3A and S3A-E). Nonetheless, putative regular spiking neurons exhibited sharper tuning to either sound-source (Figure S3F and S3G).

Notably, in a sizable fraction of neurons that classified as egocentrically untuned to either loudspeaker, the response magnitude differed significantly between the two loudspeakers (16/40, 40%, Figure 3A, “vi” in Figure 2E). Since the two loudspeakers could only be distinguished by their allocentric location, these neurons could be classified as “purely” allocentric, i.e. sound-source angle independent coding.

Response timing relative to stimulus duration is known to provide additional information about sound-source location (Middlebrooks et al., 1994; Furukawa and Middlebrooks, 2002; Mickey and Middlebrooks, 2003). We found that the spatial representations in A1 of the actively localizing gerbils also depended on the relative timing of responses, as the egocentric tuning of individual neurons frequently varied considerably across response epochs (Figures 3C and 3D). Additionally, often this change of preferred egocentric location across response time occurred only for one loudspeaker.

To test to what extent the diverse activity patterns that we observed may facilitate distinguishing the two sound-sources during task performance, we implemented an artificial neural network model with one hidden layer (multilayer perceptron classifier, see Methods). The algorithm was trained on 224 units (141 single neurons and 83 multi-units) from two animals from a total of 21 sessions and the training classes were a combination of egocentric spatial information and allocentric sound-source identity (using random under-sampling to prevent class imbalance), in which the 360° around the animal were divided in 8 classes for each loudspeaker. The neural network was then presented with test data (see Methods) for decoding of specific spatial information. Initially, the data from both loudspeakers was merged to calculate the accuracy of the decoder on the egocentric location of the active loudspeaker. The neural network model classified the egocentric locations with high accuracy (Figure 4A). Next, we determined the capability of the algorithm in identifying the active loudspeaker irrespective of the egocentric location. Remarkably, the decoding accuracy for identifying the sound-source was also highly significant (Figure 4B), demonstrating the coexistence of both subject-based and world-based reference frames in the neuronal representations (Figure 4C). Temporal analysis of model performance indicates that the egocentric position of the active loudspeaker was largely encoded within the first 100 ms after sound onset, whereas the information about the identity of the loudspeaker increased monotonically throughout a stimulation period (Figure 4D) and reached behavioral performance levels (Figure 1D).

**Figure 4.**
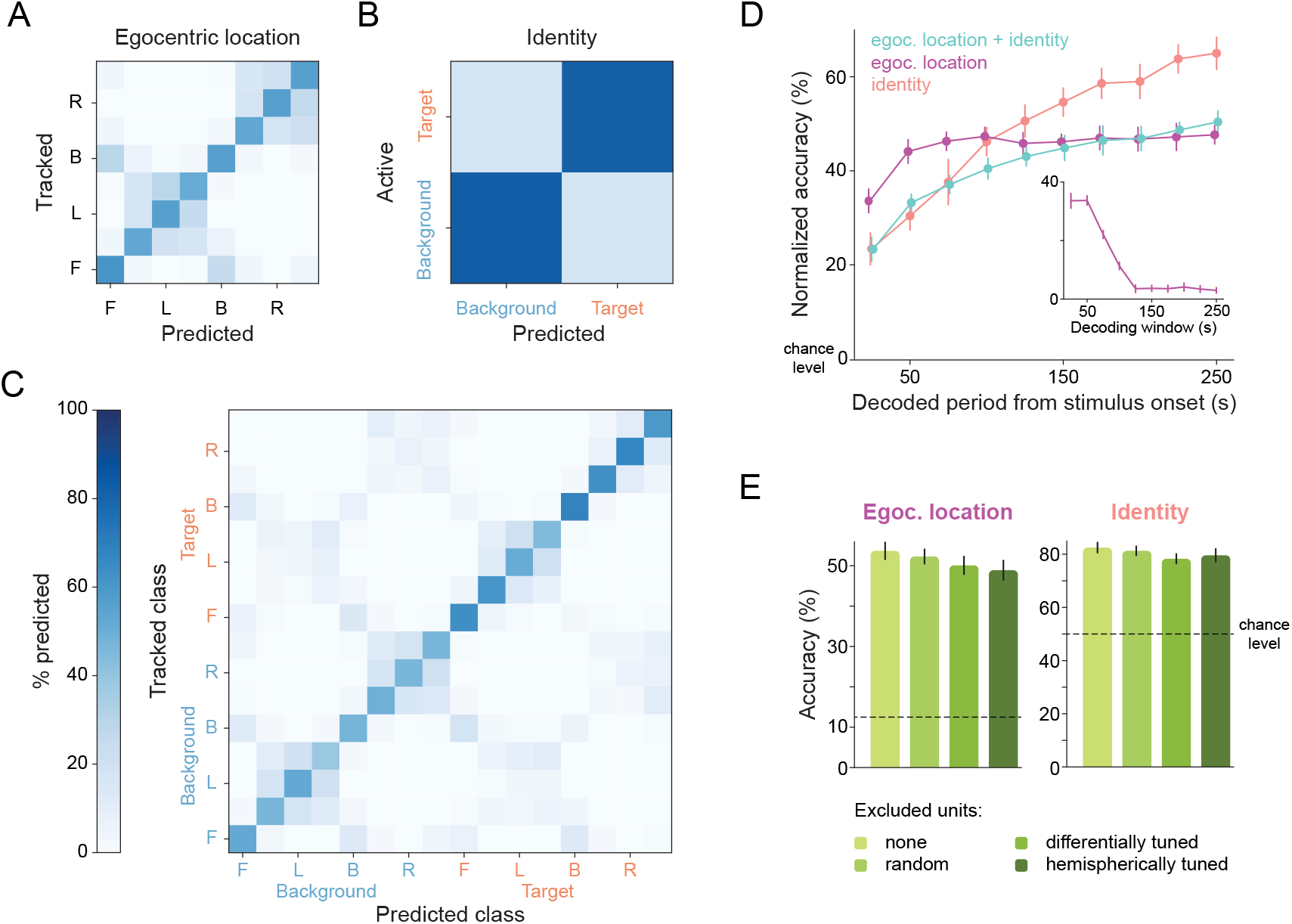
Artificial neural network decoding of population activity. Confusion matrix for decoding: (A) the egocentric loudspeaker location; (B) the identity of the active loudspeaker; (C) the combination of the loudspeaker location and identity. F: front, L: left, B: back, R: right. For the construction of all the confusion matrices, 38980 values were used across 20 sampling cycles (1949 values per cycle). (D) Relationship between the accuracy of the decoder and the time after sound onset used for decoding. Inset: Accuracy of decoding the egocentric location when using exactly one 25 ms bin. The normalized accuracy drops linearly from over 30 % until 50 ms to almost chance level after 125 ms after sound onset. All error bars correspond to the standard deviation of the mean normalized accuracy across 20 sampling cycles. (E) Eliminating 18 units of different tuning types from the test dataset resulted in a minor drop in the decoding accuracy and remained highly significant above chance (differentially tuned: −3.6 ± 2.2 % and −4.2 ± 1.8 % for location and identity, respectively; randomly selected, −1.4± 1.8% and −1.2± 1.7%, hemispherically tuned: −4.8 ± 2.4% and −2.9 ± 2.4%). All error bars correspond to the standard deviation of the mean accuracy across 100 sampling cycles.

The contribution of the diverse response patterns across the population of neurons to the egocentric and identity decoding accuracy was explored by selectively eliminating units with specific tuning characteristics in the test data and re-assessing the decoding accuracy (Figure 4E). Elimination of either all units with differential spatial tuning for the two sound-sources (18 units, compare Figure 2E, type “v”) or exclusion of the same number of units with either diverse response profiles (randomly selected) or exclusively canonic hemispherically tuned units (compare Figure 2E, type “i” and “iii”) resulted in only mild reductions of accuracy for location and identity, respectively. Overall, the neural network algorithm demonstrated remarkable robustness towards these interventions, as decoding accuracy for both reference frames remained far above chance level regardless of the eliminated neuronal tuning class (Figure S4). This robustness suggests that distributed interactions in population responses underlie an interlaced coding of egocentric and allocentric information.

## Discussion

Here, we demonstrate new cortical computations for active sensory object identification during unrestricted navigation. By introducing differential behavioral meaning to two loudspeakers at distinct locations, we find spatial representations in A1 neurons that were so far unreported.

Maybe most intriguingly, we observed that the identity of the active sound-source altered the spatial preference for physically identical stimuli. Specifically, neurons in the engaged A1 exhibit spatial tuning that combine ego- and allocentric (i.e source identity) components to various degrees. We also found a small subset of neurons whose response was modulated by the identity of the active sound-source, but that were insensitive to egocentric locations. Since in our paradigm, source identity was distinguishable only by their respective allocentric position, these neurons can be classified as purely allocentric. An earlier study (Town et al., 2017) had also reported on a comparably small subset (6.5 %) of A1 units with allocentric tuning in freely exploring but acoustically unengaged ferrets. In stark contrast to our results, however, all units in their study were classified as either exclusively ego- or allocentrically tuned.

Sound-source identity also influenced the spatial tuning acuity, which was higher to the target compared to the background loudspeaker. Sharpening of spatial tuning in A1 has been previously reported for cats when engaged in a sound localization task compared to a periodicity task (Lee and Middlebrooks, 2011), suggesting that our gerbils’ localization behavior might have changed after entering the target island.

Differential tuning to the two loudspeakers was especially apparent at later response periods (>~130 ms) which, interestingly, have previously been implicated to be crucial for the creation of auditory objects (Nelken et al., 2003) and to encode perceived pitch and vowel identity in a discrimination task (Bizley et al., 2013; Town et al., 2018). Although we did not directly probe the influence by other areas, the richness of our recorded late responses are indicative of top-down modulation (Burgess, 2008; Yin et al., 2020). Potentially relevant feedback connections originate in the retrosplenial cortex, which is strongly connected to the hippocampus and projects to AC (Budinger and Scheich, 2009) and has been suggested to transform egocentric into allocentric information (Vann et al., 2009). Allocentric representations were also found to be present in the cingulate cortex (Dean and Platt, 2006), which also projects to the auditory cortex (Budinger and Scheich, 2009).

Irrespective of their spatial tuning, the neuronal response magnitude of A1 neurons differed based on the choice of the animals to remain in the island or to leave before receiving any reward. This finding is in line with earlier studies that found A1 neurons to encode task choice (Selezneva et al., 2006; Francis et al., 2018; Guo et al., 2019), and could also be associated with predictive processing (Keller and Mrsic-Flogel, 2018) and reward expectancy (Poort et al., 2015). Ventral tegmental area (VTA) is a possible source for such reward signaling inputs via direct dopaminergic connections to infragranular layers of A1 (Brunk et al., 2019). Such dopaminergic neurons have been shown to encode reward probability, and their response magnitude to predict changes in behaviour (Babayan et al., 2018), which might be the mechanism underlying the decision of staying or leaving in the target island in our paradigm. Moreover, prefrontal cortex (PFC) has also been associated with encoding reward expectations (Rouault et al., 2019), as well as sound categorization (Jiang et al., 2018; Fritz et al., 2010). Direct feedback in rodents from PFC to both AC (Budinger and Scheich, 2009) and the midbrain (Olthof et al., 2019), renders this area also a likely candidate for top-down modulation in the computations described herein.

Together, these novel neuronal computations appear well-suited for identifying and tracking specific sound-sources in the environment during self-motion. We therefore propose that experiments in freely-moving and engaged animals are crucial to better understand long-standing phenomena of active sensing in every-day life such as the cocktail party problem (Bizley and Cohen, 2013).

## Methods

### Animals

All procedures were approved in accordance with the stipulations of the German animal welfare law (AZ 55.2-1-54-2532-74-2016). Experiments were conducted on 11 male Mongolian gerbils (*Meriones unguiculatus*) from the breeding colony of the Biocenter of the Ludwig-Maximilians University Munich. Animals were housed in groups of 3 to 4 individuals at a temperature of 22.4^°^C and 66 % humidity with 12 h light/dark cycles. The experiments were conducted during the light phase of the cycle.

### Behavioral training and stimulus control

Behavioral data was collected from eleven gerbils. Animals were required to be at least 8 weeks old at the beginning of training and underwent a general habituation period in the SIT setup for 15 minutes per day for 5 days. Gerbils had unrestricted access to water in their home cages. Food was provided as pellets ad libitum until the training phase started, after which animals were only allowed food as rewards for correct trials (half of a sunflower seed or 20 mg, TestDiet LabTab AIN-76A). The weight of every gerbil was measured daily and additional food was provided if needed to ensure the weight not to drop more than 5 % between consecutive training days and to maintain it within the desired range of 60–80 g.

Animals were trained using the SIT paradigm (Ferreiro et al., 2020) for probing sensory perception in unrestrained and actively engaged animals. In SIT, a freely moving animal was trained to search for an unknown area in the arena (target island), which prompted a change in the identity of the loudspeaker (from “background” to “target”) from which a stimulation is played back, and to report the detection of the change by remaining in the target island for a determined time interval (“sit-time” = 6 s). The position of the target island was random and changed every trial (pseudo-randomly chosen from a uniform distribution), making the loudspeaker change the only useful cue to find the island. After correctly reporting the detection of the target stimulus, the animal was rewarded with food that automatically dropped in the arena from an overhead food dispenser. Trials had a time limit of 60 s. If the animal did not correctly report the target island within the time limit, a low-pass filtered noise was presented to the animal for 10 s, during which no new trial could be initiated.

Details of the setup and stimulation control were reported elsewhere (Ferreiro et al., 2020) and are summarized here: The setup consisted of a circular arena (diameter = 92 cm) within a sound attenuated chamber (Figure 1A). Stimuli were computer generated and transmitted through an amplifier (AVR 445 Harman/Kardon, Germany) and small loudspeakers (Aurasound NSW1-205-8A 1” Extended Range) mounted externally of the arena (~5 cm distance to the perforated metal walls of the arena). Stimuli were 57 ms long harmonic complex sounds with a fundamental frequency of 147 *pm* 4 Hz and low-pass filtered below 1.5 kHz. Animals were trained to initiate a trial by remaining on an initiation platform (approx. 1cm in height, 12 cm diameter) for 1s. Trial initiation triggered the playback from the background loudspeaker (located either east or west of the initiation platform), and animal entrance into the island triggered the switch of the playback to the target loudspeaker (separated by 180^°^ from the background loudspeaker). Stimuli were played at a repetition rate of 4 Hz and their amplitude was 70 dB SPL roved ±5 dB. The animal’s position was tracked via images captured every 250 ms with a Flea3 camera (FL3-U3-13Y3M-C, Point Grey Research Inc.). Stimulation parameters (loudspeaker identity) were updated online according to the animals’ position within the arena. Custom-made software for animal tracking, stimuli generation and food reward delivery was developed in MATLAB.

No more than two training sessions were carried out per day, lasting up to 90 minutes. The training of the animals was performed by gradually reducing island size (starting at diameter = 42 cm, 21 % of the arena surface) and increasing sit-time (starting with 2 s) over the course of the training sessions. The final parameters of the island size were diameter = 25 cm (~7% of the arena surface) and sit-time = 6 s.

### Video tracking

Custom-made online tracking scripts were developed in MATLAB. In the animals’ tracking algorithm, an ellipse was fitted to the animal after background subtraction and the centroid of this ellipse was taken as the position of the gerbil. The recorded videos were offline analyzed in Python (with OpenCV library - Bradski, 2000) where the ellipse and its centroid position was re-analyzed and the orientation of the fitted ellipse determined. The median orientation error was assessed blindly to be 6^°^, with 75% of the orientation errors being below 15°.

### Behavioral data analysis

Data analyses were performed in MATLAB (Mathworks) and Python using custom scripts. To test the performance of the animals, we compared the percentage of correct trials in each session with surrogate runs based on random target island shuffling. That is, for each trial (offline, a posteriori), 1000 surrogate (non-real) islands, non-overlapping with the target one, were randomly set and the real trajectory of the animal was used to calculate in how many of these islands the trial would have been correct at each time point given the required sit-time. The median chance performance and 95% confidence interval was calculated based on bootstrapping (random sampling with replacement from all the trials of the session). This method allows obtaining an estimate of the proportion of correct trials the animal would have gotten just by chance given their locomotion trajectory and dynamics.

### Surgery

Electrophysiological data was collected from five trained gerbils. Animals were anesthetized with an intraperitoneal injection of a mixture of metedomidin (0.15 mg/kg), midazolam (7.5 mg/kg), and fentanyl (0.03 mg/kg). To maintain it at a constant level, the same mixture was subcutaneously re-injected every 90 min. After shaving and disinfecting the head, a local anesthetic (50 μl, 2 % xylocaine) was injected under the scalp skin and below the skin near the ears. For protection and to prevent dehydration, the eyes were covered with an ophthalmic gel (Thilo-Tears SE, Alcon Pharma Gmbh). The animal was then transferred to the stereotactic apparatus, where its head was securely fixed via a bite and ear bars. Its internal temperature was monitored with a rectal thermometer and kept constant at 37^°^C throughout the experiment by a feedback controlled electric heating pad (Harvard Apparatus). After disinfection, a midline scalp incision was performed to expose the skull. Subsequently, the connective tissue on the skull was removed with a bone curette and the skull was treated with 35 % phosphoric acid (iBOND etch gel, Kulzer), which was promptly washed away. Structural screws were placed on top of the left frontal and right parietal bones and the ground screw on the occipital bone, so that it gently touched the brain. After stereotactic alignment, a 3×3 mm craniotomy and durotomy were performed on top of the left auditory cortex, followed by a very slow lowering (2 μm/s) of a tetrode bundle to a maximum depth of 0.9 mm into the cortex, using a micromanipulator (Scientifica). The craniotomy was carefully filled with KY-jelly and immediately sealed with dental cement (Paladur, Kulzer), which also fixated the bottom of the microdrive and the outer cannula that protected the tetrodes. 1 ml of Ringer’s solution was subcutaneously injected at the end of the surgery and the anesthesia was reversed via subcutaneous injection of the antagonist mixture composed of naloxone (0.5mg/kg), flumazenil (0.4mg/kg), and atipamezol (0.375mg/kg). Analgesics (0.2mg/kg, meloxicam) and antibiotics (7.5 mg/kg, enrofloxacin) were orally administered post surgically for five subsequent recovery days. During this time, the animals had food and water ad libitum and were not trained.

The implant used in this experiment was a tetrode bundle consisting of four tetrodes glued together, which, on their turn, consisted of four insulated tungsten wires (12.7 *μm* diameter each, tungsten 99.95%, California Fine Wire) twisted around each other. Each wire was connected to a custom-made printed circuit board with Omnetics connector (Axona), which was attached to a lightweight microdrive (0.25 mm/turn, Axona). The tetrodes were glued together and protected by an inner and outer cannula that could slide by each other. On the day prior to the surgery, the tip of all electrodes were cut with sharp scissors and gold plated (Non-Cyanide Gold Plating Solution, Neuralynx) to reach a desired impedance of 100-150 kΩ (at 1kHz). The tetrode bundle was implanted vertically in the following coordinates from lambda: 6.2 mm lateral, 2.6 mm anterior.

### Electrophysiological recordings during task performance

Recordings were made via a wireless headstage (W2100-HS16, Multichannel Systems). The physiological signals were at first amplified between 1 Hz and 5 kHz and digitized (16-bit resolution) in the headstage, then wirelessly transmitted to the receiver (W2100-RE-AO, Multichannel Systems) at a sampling rate of 25 kHz and recorded by a PC via an interface board (MCS-IFB 3.0 Multi-boot, Multichannel Systems), and commercial software (Multi Channel Experimenter, Multichannel Systems). Simultaneous with the onset of sound presentation, a short signal was sent via sound card to the analog input of the interface board, allowing the synchronization of the physiological recordings with the sound presentation, and a digital signal was transmitted to the interface board indicating the beginning and end of the trial, which was later used to align the video information.

### Spike sorting

Initially, the raw electrophysiological signals were high-pass filtered above 300 Hz and a common median referencing (Rolston et al., 2009) was performed for the whitening among all channels and removal of large artifacts. The signals were low-pass filtered below 5 kHz and fed to a spike sorting algorithm based on template matching (Kilosort - Pachitariu et al., 2016). Afterwards, the automatically sorted spikes were manually inspected and the corresponding clusters refined with the graphical user interface phy (Rossant et al., 2016). Only units with an isolation distance larger than 20, more than 200 spikes and less than 2 % of the spikes within the 2 ms refractory period were considered single cells (Schmitzer-Torbert et al., 2005; Hill et al., 2011). All units that passed the manual curation but not the isolation distance or refractory period requirements were considered multiunits.

### Peristimulus time histograms and response epoch analysis

The peristimulus time histogram (PSTH) was calculated based on bootstrapping data. An algorithm sampled 500 times (with replacement) from the total number of recordings of a given sound. For each cycle, a histogram was calculated with binning width 8 ms and the median and 95 % confidence intervals were determined for the total of bootstrapping cycles normalized by the total number of analyzed sounds. For the calculation of neuronal latencies without contamination by baseline activity, the Bayesian blocks method (Scargle et al., 2013) was used, which identifies statistically significant variation in a time series to optimally segment the data. If at least one new edge is created, the unit is considered modulated and therefore auditory responsive. The latency of a given unit was defined as the first created edge. To determine whether the onset response corresponds to an increase in the firing rate, the maximum displacement of the median values of the firing rate in the 15 ms after latency was compared to the baseline (last 50ms before latency). If it was lower than the mean of the lower bound of the baseline’s 95 % confidence interval, the onset firing rate was considered to be decreasing and if it was higher than the mean of the upper bound of the baseline’s 95 % confidence interval, the onset firing rate was considered to be increasing. The same Bayesian blocks method was employed to calculate whether a unit had a significant offset response within the interval [latency+50:latency+70] ms. If so, the offset period was defined as the time point of the new edge creation, otherwise it was defined as latency + sound duration (57 ms). The start of the late response period was defined as the start of the offset period + sound duration (57 ms).

Mean firing rates were calculated as the mean of the PSTH across all time bins.

For the comparison between the first and last 2 s inside the island and whenever the animal wrongly left it, only situations in which the animal stayed at least 2 s in the island were considered. Furthermore, if the last 2 s overlapped in time with the first 2 s (the animal stayed less than 4 s in the island), only the non-overlapping part was used for the calculation of the last 2 s.

### Spatial tuning analysis

For the spatial tuning analysis, the 360^°^ angular space around the animal was divided in 8 bins. A minimum of 10 sound presentations per binned angle was a primary condition for the spatial tuning analysis, as well as a maximum difference of 10 between the bins with the most and the least number of sound presentations. A 1000 cycle bootstrapping method with replacement was implemented to calculate the spatial tuning, in which an angular histogram was calculated for the corresponding spikes and normalized by the number of sounds at each egocentric sound-source angle bin for the chosen loudspeaker. The vector strength and corresponding preferred egocentric sound-source angle were then calculated for each bootstrapping cycle. As some units revealed a more complex spatial tuning (with two peaks in opposite directions), the folded vector strength (in which responses to angles opposite to each other are summed - Mazurek et al., 2014) was additionally calculated together with the corresponding orientation angle.

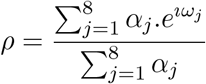

where 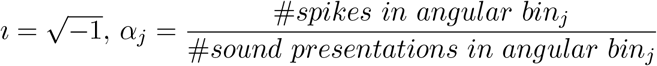 and 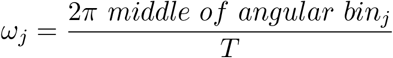.

The vector strength is *VS* = |ρ| with *T* = 360 and the corresponding preferred egocentric sound-source angle *ang_dir_* = arg(ρ). The folded vector strength is *VS_folded_* = |ρ| with *T* = 180 and the corresponding preferred egocentric sound-source angle angle *aug_ori_* = arg(ρ)/2. From the bootstrapped data, the 95% confidence interval for each angle bin was calculated, as well as the 68 % confidence interval of the vector strength and orientation vector. A unit was considered spatially tuned at a particular temporal period (onset, offset or late) if the vector strength (or folded vector strength, whichever is larger) was larger than 0.2 and the lower bound of 68% confidence interval of the vector strength was larger than 0.15.

Units were considered as egocentrically untuned if the median vector strength was smaller than 0.15 in all three phases of the PSTH for both loudspeakers. The firing rate in response to the two loudspeakers was considered significantly different if there was no overlap in the 95 % confidence interval of the calculation of the mean firing rate for each loudspeaker.

### Decoder Analysis

For the decoder analysis, the decoder was trained on the recorded responses of 224 units (141 single neurons and 83 multi-units) from two animals in a total of 21 sessions. Each sound presentation was categorized with respect to the active loudspeaker and the egocentric sound-source angle at the time of sound onset (in 8 angular bins). Therefore, each sound presentation corresponded to one of the 16 possible loudspeaker-angular bin classes. Spike counts, for each recorded unit, in each of the 250 ms response periods were determined in 10 bins, each with a 25 ms duration, and then normalized between 0 and 1 (using the mix-max scaler algorithm) to avoid that units that are inherently more active dominate the results. These unit responses were then pooled across sessions for each of the angular-bin classes, creating 16 across-sessions class-specific population-response profile, which were then used to train the decoder. To prevent class imbalance, due to non-uniform occurrence of angular-bin classes during the task, random under-sampling (without replacement) was implemented to feed the decoder with the same amount of data per class. The test dataset consisted of a minimum of 25 % of the population responses per class.

The training data was fitted by a multi-layer perceptron classifier (MLPC - Pedregosa et al., 2011), a feedforward artificial neural network which uses backpropagation during training for optimization of the parameters in a supervised learning manner, and therefore was interpreted as a more biologically inspired decoder. In the MLPC, one hidden layer was implemented with the number of nodes as the mean between the number of features used and the total number of classes (16). The optimization algorithm used was the “lbfgs” from the family of quasi-Newton methods and the biologically-inspired rectified linear unit was the activation function for the hidden layer.

The whole process was repeated 20 times to estimate errors (100 times in the case of the unit-elimination analysis). The accuracy of identifying the active loudspeaker was determined per loudspeaker for each sampling repetition, later the accuracy per repetition was considered as the mean between the two loudspeakers and the total accuracy and standard deviation calculated across all 20 sampling turns. The accuracy of predicting the right class and egocentric sound-source location was similarly calculated. For the construction of the confusion matrices, all the predictions from all the sampling repetitions were simultaneously used.

To evaluate the influence of units with different spatial tuning, units were eliminated in the test data by setting all the bins corresponding to the chosen units to zero. The accuracy of the decoder (trained on the training data from all the units) was then compared to when the test data was complete. In the elimination of the random units, different units were chosen to be eliminated in each sampling turn. The units with differential spatial tuning for both loudspeakers which were eliminated corresponded to all the units in the training set which at some temporal response period had a difference in preferred egocentric sound-source angle between loudspeakers larger than 80^°^. The eliminated canonical units were randomly selected from the neurons which showed clear ipsi or contralateral tuning during the onset period.

### Histology

After a 350 *μ*l intraperitoneal injection of pentobarbital sodium, the animal was perfused with 4 % paraformaldehyde (PFA) and the brain carefully removed and stored in PFA in the fridge. Afterwards the brain was twice washed in PBS and the frontal part cut with a vibrotome in 70*μ*m thick slices. The slices were stained with green fluorescent Nissl (NeuroTrace™ 500/525) and then compared to the gerbil brain atlas (Radtke-Schuller et al., 2016) for determining the location of the recording sites.

### Statistics and reproducibility

All error bars correspond to the standard deviation unless stated otherwise. All shaded areas correspond to the 95 % confidence interval as calculated via bootstrapping. For comparisons of central tendencies on the group level, we used two-tailed non-parametric tests: Wilcoxon signed-rank tests for paired samples and Mann-Whitney U-test for independent samples. All hypotheses were tested at an alpha level of 0.05.

Data collection and analysis were not performed blind to the conditions of the experiments. Our experimental design provided a within-animal control, and because comparisons were not required between different groups, blinding was not necessary. All analyses were based on automated scripts applied across animals and thus were not subject to any experimenter bias.

### Data and code availability

At the time of publishing, the data and code used in this study will be available upon reasonable request from the lead contact.

## Acknowledgements

This study was supported by the Deutsche Forschungsgemeinschaft DFG (PE2251/2-1 to MP, and SFB 870, project B02 to MP and BG), and by the International Max Planck Research School for Molecular Life Sciences (to DA). We thank N. Lesica and P. Alcami for valuable comments on earlier versions of the manuscript and H. Wohlfrom, O. Alexandrova and K. Fischer for assistance with the histology. We are grateful to L. Wiegrebe for help on the behavioral paradigm.

## Author Contributions

MP, DA, and BG conceived the study. MP and DA designed the experiments and DNF and BG contributed to paradigm refinement. DA performed the experiments and analyzed the data. DA, MP, and DNF designed the data presentation and wrote the manuscript. All authors provided comments and approved the manuscript.

## Supplemental information

**Figure S1.**
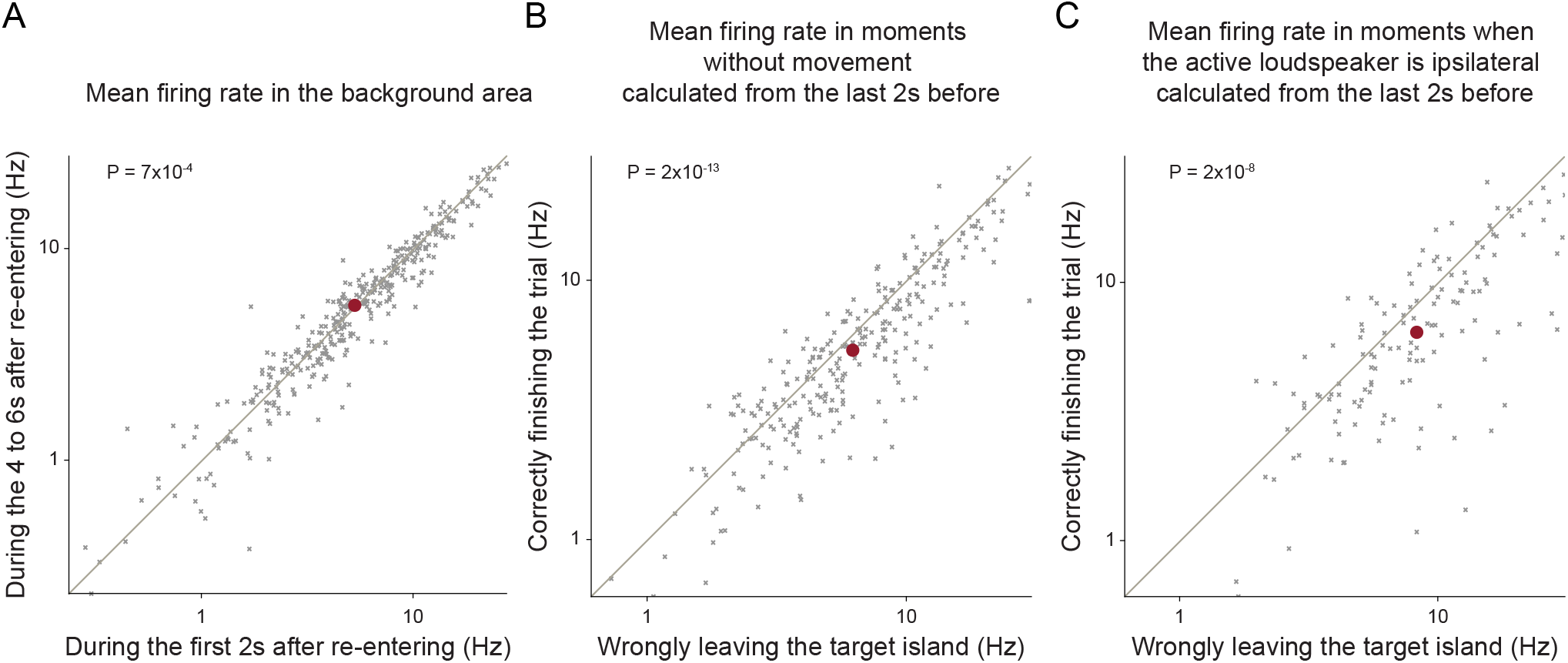
Changes in firing rate are specific to trial-outcome. The decrease in mean firing rate when the animal finishes the trial is not due to: (A) unspecific neuronal rate adaptation, as no firing rate decrease is observed in the initial 6 s of background stimulation (360 neurons); (B) movement, since the reduction in firing rate persists even when calculated exclusively for periods without movement (267 neurons); (C) a spatial bias in the orientation of the animal, because the decrease persists even when calculated exclusively for periods when the active loudspeaker was exclusively ipsilaterally located (137 neurons). The Wilcoxon signed-rank test was used in all analysis.

**Figure S2.**
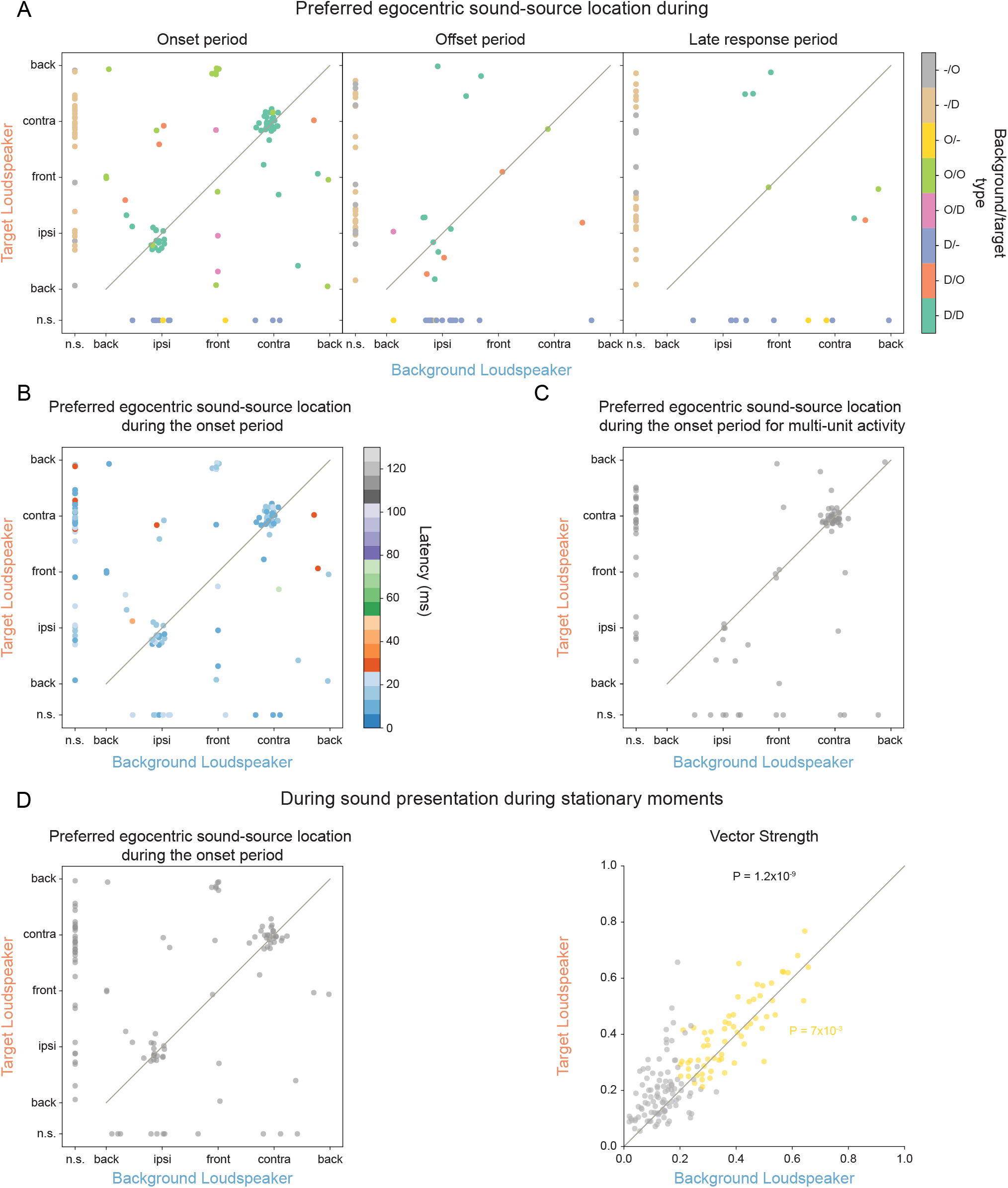
Preferred egocentric sound-source location correspondence between the target and the background loudspeakers. (A) during the onset (left), offset (center) and late response periods (right) color-coded with the type of spatial tuning for each loudspeaker; D: direction-sensitive O: orientation-sensitive; (B) during the onset period, color-coded with the latency determined for each neuron; (C) during the onset period, for the multi-unit activity; (D) during stationary moments. (E) Vector strength comparison between target and background during stationary moments.

**Figure S3.**
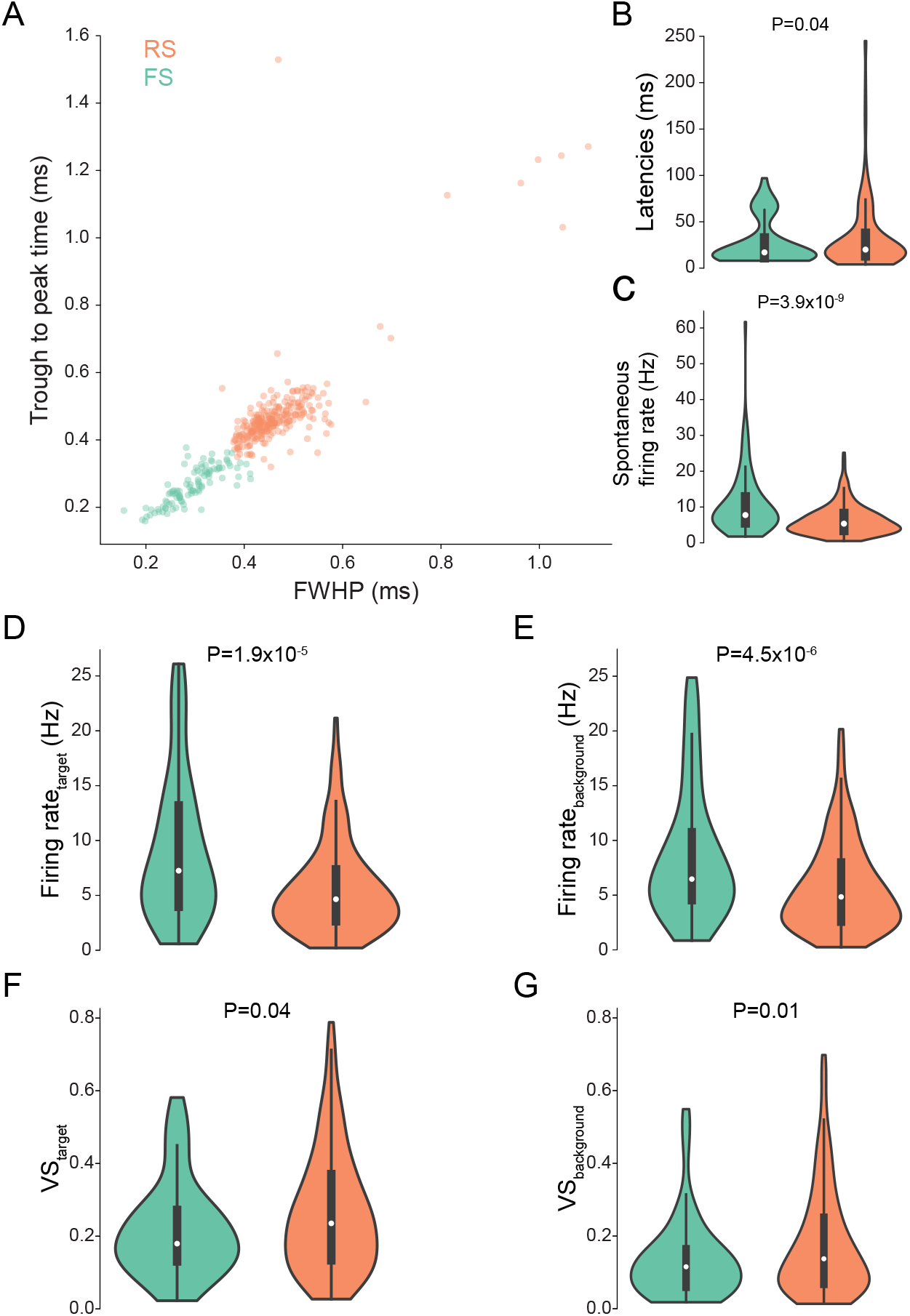
Different response properties of neuronal types. (A) Response properties of putative regular spiking (RS) and fast spiking (FS) neurons were separated based on spike-waveform properties by k-means clustering analysis. The clustering was performed on the full width at half maximum of the peak (FWHP) and of the trough, and on the trough to peak time. (364 neurons: 102 FS and 262 RS). On the population level, FS neurons exhibited: (B) shorter latencies (364 neurons); (C) and higher mean firing rates, both for spontaneous activity (364 neurons); (D) as well as in response to the target (244 neurons); (E) or background loudspeaker (333 neurons). (F) and (G) Spatial tuning was sharper in RS neurons (204 and 354 neurons for target and background, respectively). All p-values based on the Mann-Whitney U-test.

**Figure S4.**
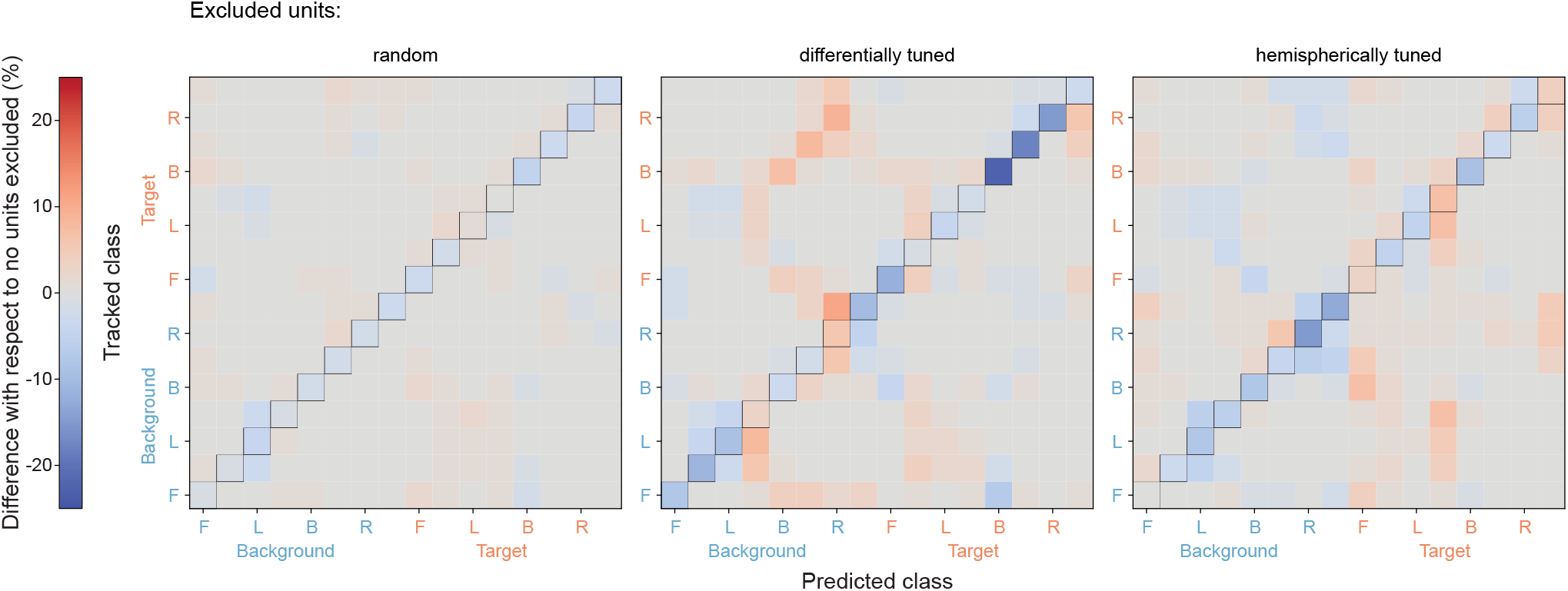
Influence of eliminating neuronal tuning classes on the decoding accuracy. Difference in the confusion matrix relative to the original test dataset when 18 randomly selected units (left), 18 units with differential spatial tuning (middle) or 18 canonic hemispherically tuned units (right) are excluded from the test dataset (see also Figure 4E). Number of sampling cycles: 100.

**Video S1. Example trial of an animal performing in the task**

Left: Allocentric perspective: The animal starts a trial by staying 1 s on top of the initiation platform (yellow circle) while its position and orientation is tracked (green ellipse and arrow). The moments of; sound stimulation are represented by the blinking blue circle (background loudspeaker) or the orange circle (target loudspeaker. After staying 6s in the island (orange circle), a reward is dropped. Right: Egocentric perspective: Representation of the location of the active loudspeaker relative to the animal - blue if the background loudspeaker is active and orange if the target loudspeaker is active.

